## Supplementary Figures for "Isobaric crosslinking mass spectrometry technology for studying conformational and structural changes in proteins and complexes"

401 Terry Avenue North

Seattle, WA 98109

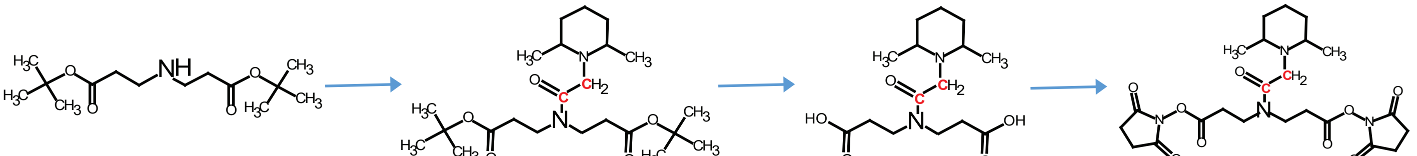

1 2 3 4

1. 1-$^{13}$C or 2-$^{13}$C BAA /DIC
2. 2,6-dimethylpiperidine
3. RP-C18

1. TFA
2. RP-C18

1. TSTU/DIPEA

**Figure S1. Synthesis of Q2linkers.** The $^{13}$C atom in C1q2 or C2q2 is indicated by red font.

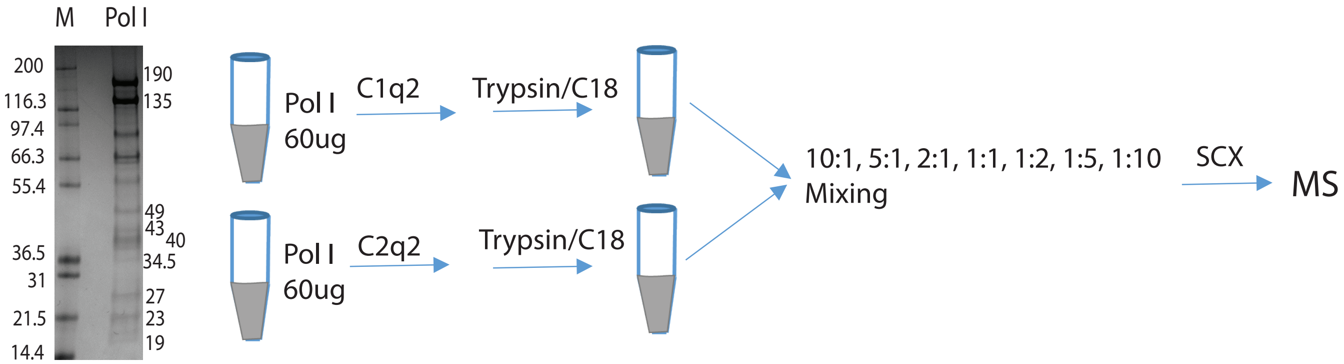


**Figure S2**. **Experimental procedure to evaluate the ability of Q2linkers to quantify the relative abundances of crosslinks and monolinks derived from Q2linker modification of affinity purified pol I.** M = molecular weight markers.

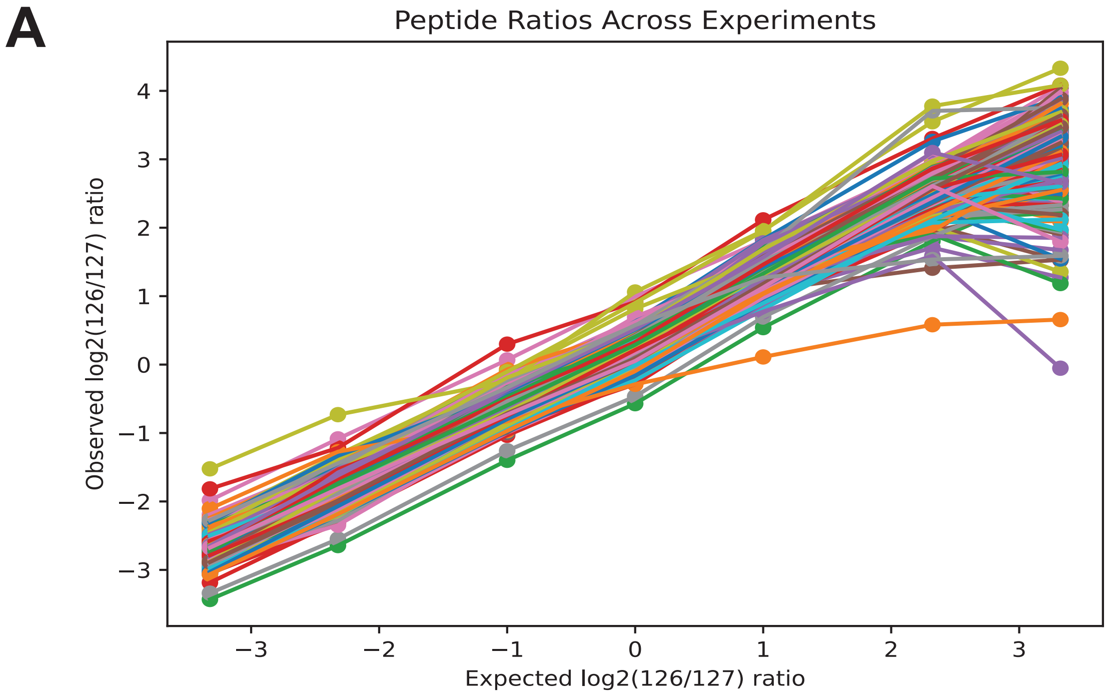


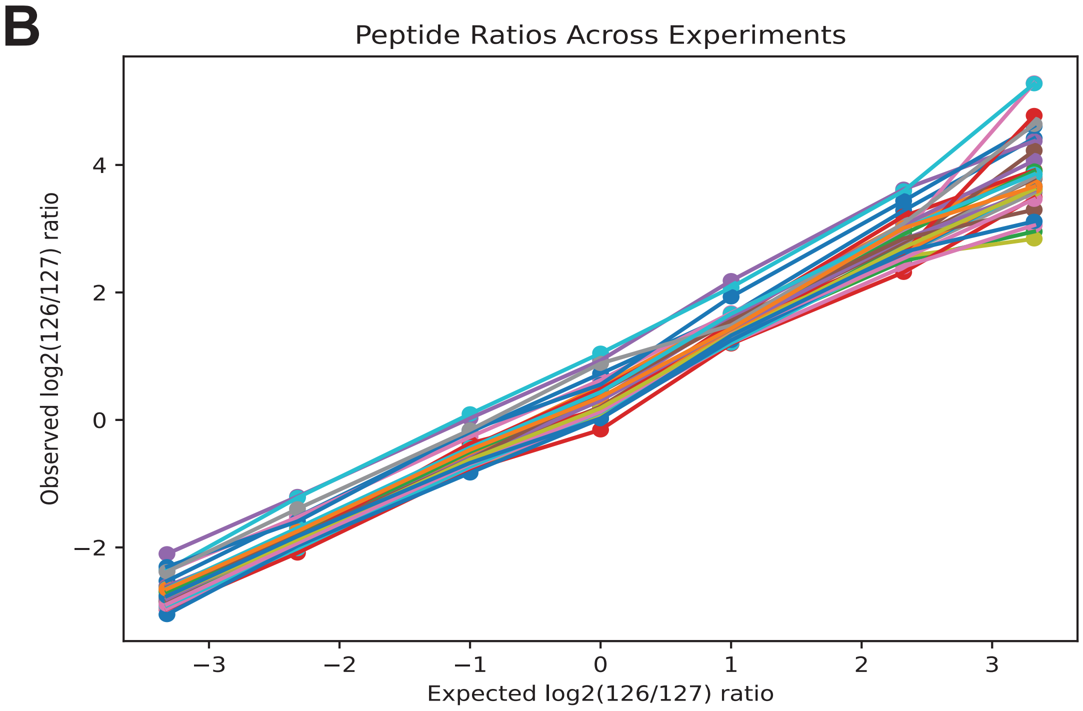


**Figure S3. Observed vs. expected log2 (126/127) reporter ion ratios for individual monolinks (A) and crosslinks (B) at different mixing ratios**. There are 298 monolinks and 32 crosslinks identified in all runs.

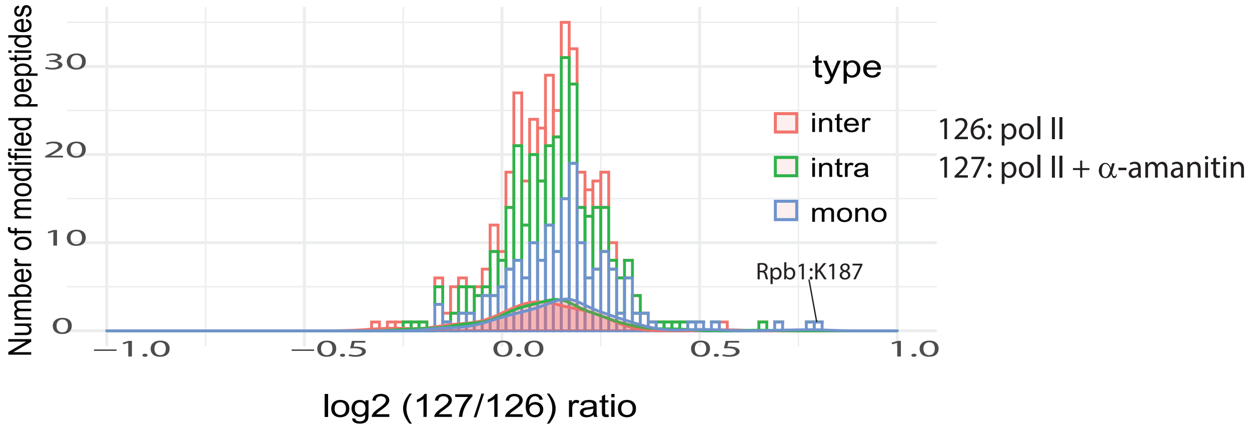


**Figure S4. The ratio distributions for interlinks, intralinks and monolinks from the pol II +/- α-amanitin experiment.**

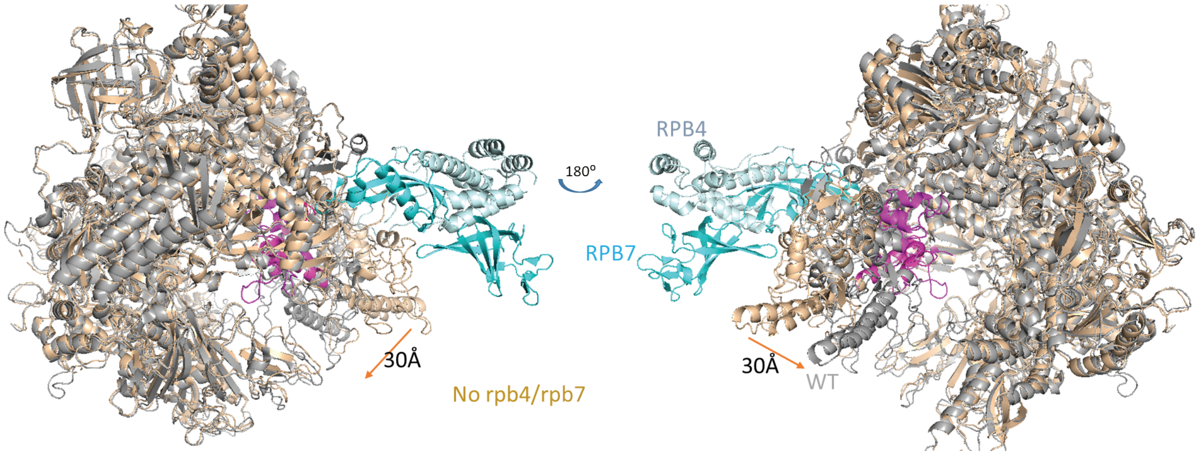


**Figure S5. Structural comparison of holo-pol II (5u5q) and core-pol II (li3q).** Core-pol II without Rpb4/Rpb7 is colored brown and holo-pol II is colored gray with Rpb4 and Rpb7 in light blue. The five switches that cause the movement of the clamp are colored in magenta.
